# Identification and Characterization of a Novel Nematode Pan Allergen (NPA) from *Wuchereria bancrofti* and their Potential Role in Human Filarial Tropical Pulmonary Eosinophilia (TPE)

**DOI:** 10.1101/2023.12.04.569914

**Authors:** Samuel Christopher Katru, Gnanasekar Munirathinam, Azadeh Hadadianpour, Anand Setty Balakrishnan, Scott A. Smith, Ramaswamy Kalyansundaram

**Affiliations:** Department of Biomedical Sciences, University of Illinois, College of Medicine Rockford, IL, United States; Vanderbilt University Medical Center, Department of Pathology, Microbiology and Immunology, A2210 Medical Center North, 1161 21st Avenue South, Nashville, Tennessee, 37232-2582

**Keywords:** Lymphatic Filariasis, Tropical Pulmonary Eosinophilia, *Wuchereria bancrofti*, Histamine, Phage display, Pan Allergen, IgE antibodies

## Abstract

Tropical pulmonary eosinophilia (TPE) is a chronic respiratory syndrome associated with Lymphatic Filariasis (LF), a tropical parasitic infection of the human, transmitted by mosquitoes. A larval form of LF, the microfilariae trapped in the lungs of TPE subjects have a major role in initiating the TPE syndrome. To date, there are no reports on the potential allergen that is responsible for generating parasite-specific IgE in TPE. In this project, we screened a cDNA expression library of the microfilarial stages of *Wuchereria bancrofti* with monoclonal IgE antibodies prepared from subjects with clinical filarial infections. Our studies identified a novel molecule that showed significant sequence similarity to an allergen. A blast analysis showed the presence of similar proteins in a number of nematodes parasites. Thus, we named the molecule as Nematode Pan Allergen (NPA). Subsequent functional analysis showed that NPA is a potent allergen that can cause release of histamine from mast cells, induce secretion of proinflammatory cytokines from alveolar macrophages and promote accumulation of eosinophils, all of which occur in TPE lungs. Therefore, we believe that NPA may have a significant role in the pathology of the TPE syndrome.

## 1 Introduction

Tropical Pulmonary Eosinophilia (TPE) is a hyper respiratory syndrome that is an occult manifestation of lymphatic filariasis (LF) infection, a filarial parasitic infection caused mainly by *Wuchereria bancrofti* (*Wb*) and *Brugia malayi* [1, 2]. According to the World Health Organization (WHO), LF infection is endemic in 52 countries in Asia, Africa, South America, and the Caribbean, with around 83 million people infected with this disease and another 886 million at risk [3]. Approximately, 1% of the LF patients show TPE symptoms [4]. If the cases are not diagnosed early and treated, the TPE syndrome can lead to severe respiratory morbidities and death [5, 6].

The LF parasites require two hosts to complete their life cycle. The insect hosts are mosquitoes belonging to the genera *Aedes, Anopheles, Mansonia*, and *Culex.* During a blood meal, the mosquitoes pick up the circulating larval stages of LF, the microfilariae, which are produced by the female worms. The microfilariae undergo development within the body of the mosquito to become the infective third stage larvae (L3). The L3s migrate to the proboscis of the mosquito and during another blood meal, the mosquito transmits the infective stage larvae into the skin of a human. The larvae migrate into the lymphatic system to develop into mature male and female worms that produce the microfilariae, completing their life cycle [7].

The microfilariae circulate in the blood during dusk, when the mosquitoes are highly active. However, during the daytime, the microfilariae settle down in various tissues including lung [8]. The metabolic products including allergens released by the microfilariae and possible allergens released from dead microfilariae can trigger an allergic immune response including the production of IgE antibodies and activation of mast cells in the lungs [9]. The inflammatory response also draws large numbers of eosinophils to the site of inflammation in the lungs leading to a complex hypersensitivity reaction that is typical of TPE [10]. Thus, accumulation of large numbers of eosinophils and neutrophils, activation of mast cells, presence of high levels of parasite-specific IgE antibodies are hallmarks of LF induced TPE in the human [4, 11]. Given its ability to activate mast cells, basophils and dendritic cells, IgE antibodies are thought to play a major role in the initiation and amplification of TPE [9, 12, 13]. Previous studies showed that a 23/25kDa protein (Bm2325) of *B. malayi* is associated with TPE [14]. However, the specific antigens responsible for the generation of IgE antibodies in TPE subjects is not fully characterized. Identifying the allergen(s) targeted by IgE antibodies in TPE might broaden our understanding on the mechanism of TPE and it will be possible to develop intervention strategies to prevent or alleviate the pathogenic symptoms of TPE [15]. Thus, the major focus of our studies was to identify and characterize the antigen(s) of the *W. bancrofti* microfilariae that binds to human IgE monoclonal antibodies (mAbs) prepared from subjects with filarial infections, some of which exhibited TPE clinical symptoms [16]. For screening the human IgE mAbs, we used a phage displayed cDNA expression library of *W. bancrofti* microfilariae as described previously [17, 18]. In this study, we report the identification of a novel nematode pan allergen (NPA) and some of the characteristics of this molecule.

## 2 Materials and methods

### 2.1 Filarial-Specific Human monoclonal IgE antibodies

IgE mAbs were generated from subjects with filarial infections, some of which exhibited TPE clinical symptoms. Preparation of the IgE mAbs are detailed in a recent publication [16]. We performed our analyses with 15 purified IgE mAbs selected from the larger panel of IgE antibodies.

### 2.2. Construction of phage library

The *Wb* mf cDNA library was obtained from Dr. Steven Williams (Smith College, Northampton, MA). For preparing the cDNA phage display expression library of *Wb* mf, we followed our previously published method [17, 19]. Briefly, aliquot of the *Wb* mf cDNA library was packaged in T7 phage library, select 10-3b packing extract (Millipore Sigma, Burlington, MA) at 22°C for 2hr and neutralized the reaction using TB (Terrific broth) medium. The packaged phages were stored at 4°C. Plaque assay was used to determine the titer of the phages. Packaged phages diluted in pre-warmed top agarose containing BLT5403 (provided with the T7 cloning kit) bacterial culture were poured onto LB (Luria-Bertani) agar plates with 50μg/ml ampicillin. The plates were incubated at 37°C until plaques were formed. The number of plaques were then manually counted.

### 2.3 Biopanning of *Wb* Mf cDNA phage expression library

For biopanning, we followed the protocol previously published by Gnanasekar et al. [17]. Pooled aliquots from the 15 IgE mAbs were used to coat the wells of a 96-well plate. The amplified phages were added to the wells, incubated and the unbound phages were washed off. The bound phages were collected using 1%SDS buffer and added to the BLT5403 culture until the cells were lysed and the collected supernatants were used in the 1^st^ round of biopanning. Plaque assay was performed using the supernatant and the number of plaques counted. This procedure was repeated for another four rounds of biopanning to enrich the phages that are specific to the pooled IgE mAbs.

### 2.4 Determining the specificity of the phage plaques to infection sera

Plaque lift assay was performed after the fifth round of biopanning on the bound phages using nitrocellulose membrane. Following this, the membrane was incubated with (i) the pooled IgE mAbs, (ii) sera samples from patients showing clinical symptoms of TPE, (iii) sera samples from patients with circulating microfilariae in their blood (MF), and (iv) sera samples from subjects who are naturally immune to the infection in the endemic areas (EN). All the human sera samples were obtained from late Dr. M.V.R. Reddy (Mahatma Gandhi Institute of Medical Sciences, Sevagram, India) with IRB approval. HRP labelled goat anti-human IgE antibodies (Invitrogen, Waltham, MA) were used as the secondary antibodies for detection and the color wasa developed using TMB substrate (Thermo Fisher Scientific, Rockford, IL).

### 2.5 PCR amplification and Bioinformatic analysis

Following biopanning, 15 phage plaques were selected and the insert DNA in the phages were PCR amplified. The PCR product obtained were Sanger sequenced at the University of Illinois DNA sequencing facility. The DNA sequence obtained were BLAST analyzed at NCBI site. With the DNA sequences obtained, a phylogenetic tree analysis was constructed to determine any evolutionary relationships with other parasitic organisms. A Clustal W analysis was performed to compare the sequences for similarity with other organisms.

### 2.6 Cloning and expression of Bm2855

Blast analysis identified Bm2855 as one of the proteins recognized by the IgE mAbs. To further characterize this protein, Bm2855 gene was amplified from the *W. bancrofti* microfilariae cDNA library. A PCR was performed using T3 vector specific forward primer (5’- AAT TAA CCC TCA CTA AAG GG) and Bm2855 gene specific reverse primer (5’- CTA TCT CGG AGC TCC TTT GGT TAC AG -3’). The parameters for PCR were 95°C of denaturation for 5 min, 95°C of denaturation for 1 min, 52°C of primer annealing for 1 min, 72°C of primer extension for 1 min and cycled for 30 cycles. A final extension of 5 min at 72°C. The PCR products were Sanger sequenced at the DNA sequencing facility of the University of Illinois.

### 2.7 Stage-specific expression of Bm2855 in *W. bancrofti*

Bm2855 was PCR amplified from the various cDNA libraries of microfilariae (MF), adult female (AF), and infective stage (L3) of *W. bancrofti* (supplied by Dr. Williams) using Bm2855 sequence-specific primers. The PCR products were separated on a 1% agarose gel and was visualized.

### 2.8 Expression and purification of rBm2855

For expression and purification of the recombinant protein, we followed the protocol already published from our lab [19]. Briefly, the recombinant plasmids were constructed in pRSET A vector, with T7 promoter, 6X His-tag, with *Bam HI*, and *Eco RI* restriction sites. The recombinant plasmid was transformed into One Shot™ BL21 Star™ (DE3) Chemically Competent *E. coli* cell (Invitrogen). When culture reached 0.6-0.8 at OD_600_, 1mM IPTG (Isopropyl β-d-1- thiogalactopyranoside, Thermo Fisher Scientific) was added to induce gene expression. Expression was checked on 12% SDS gel after staining with Coomassie blue and Western blot analysis was performed using HRP conjugated penta-His mouse mAb (Thermo Fischer Scientific). After confirmation the expression of the protein, the inclusion bodies in the cell pellets were lysed using an inclusion body solubilizing reagent (Thermo Fisher Scientific) and the supernatant was ran through an immobilized metal affinity column chromatography using HisPur™ cobalt resin (Thermo Fisher Scientific) and eluted. The proteins were desalted using Zeba spin 9k MWCO desalting column (Thermo Fisher Scientific) and concentrated using Pierce concentrators 10K MWCO (Thermo Fisher Scientific). Endotoxin free buffers and water were used in the purification steps. Endotoxin levels in the final preparation was estimated using Pierce LAL Chromogenic Endotoxin Quantitation Kit (Thermo Fisher Scientific). Finally, any trace amounts of endotoxin if any was removed using ToxinEraser™ Endotoxin Removal Kit (Thermo Fisher Scientific). This ensured that the final product has no endotoxin.

### 2.9 Specificity of monoclonal IgE antibodies

For biopanning, we used pooled IgE mAbs. Since we did not know which of the IgE mAbs bound to Bm2855, in these studies we screened each of the IgE mAbs for their binding to rBm2855. We used an indirect ELISA to measure binding. Approximately 1 ug of the rBm2855 was coated on to ELISA plates. After blocking with 5% BSA, each of the IgE mAbs (at 1:100 dilution) were added as the primary antibodies and HRP-labelled goat anti-human IgE antibody (at 1:1,000 dilution, Invitrogen, Waltham, MA) was used as the secondary antibodies. Color was developed using TMB substrate (Thermo Fisher Scientific) and OD read at 450 nm in an ELISA reader (BioTek Synergy2, Santa Clara, CA).

### 2.10 Histamine release assay

Peritoneal cells were collected from four male Balb/c mice by a peritoneal lavage using complete RPMI medium and cells isolated by centrifugation at 200g. Use and care of animals in these studies were approved by the IACUC committee of the University of Illinois Rockford. Percentage of mast cells among the peritoneal cell population was determined after staining an aliquot of cells with Hema3 kit (Fisher Scientific).

Approximately, 6×10^5^ mast cells/ml were seeded into the wells of a six well plate and incubated with 1 µg of the endotoxin-free rBm2855. The plates were incubated at 37°C for 30 mins. The amount of histamine released into the culture supernatant was determined using a histamine detection kit (Abcam, Cambridge, MA) as described previously [19]. After adding the histamine reaction buffers, samples were incubated at 37°C for 30 min and the absorbance was read at OD_450nm_. Data were analyzed after constructing a standard curve and the trend line equation as per the instructions from the manufacturer.

### 2.11 Release of proinflammatory cytokines from mouse alveolar macrophages

Alveolar macrophages were collected from the lungs of four Balb/C mice as described previously [20, 21]. After counting the cells, 3×10^6^ cells were added to the wells of a 48 well culture plate. About 1µg of the endotoxin-free Bm2855 protein was added to the cells and incubated for 48 hours at 37^0^C. Following incubation, the culture supernatant was collected. A Th1/Th2/Th17 Cytometric Bead Array kit (BD Bioscience, Franklin Lakes, NJ) was used to detect the presence of secreted cytokines. Data was acquired and analyzed in a BD FACS Calibur (BD Biosciences) flow cytometer using FlowJo software.

### 2.12 Evaluating eosinophil migration into the peritoneal cavity of mice

In these experiments, we evaluated the ability of rBm2855 to promote eosinophil accumulation in the peritoneal cavity of ovalbumin-sensitized mice as described previously [22]. Briefly, eight Balb/c mice were primed i/p twice with 20µg of ovalbumin. One week after priming, four mice were challenged i/p with 5µg of endotoxin-free rBm2855 and four control mice received endotoxin-free saline. The following day, peritoneal cells were collected by a peritoneum wash and cytospin (Wescor, CytoPro, Oak brook, IL) preps of the cells were stained with Hema3 staining kit. The number of each cell population in 10 consecutive microscopic fields were counted under 400 x. Percent cell population and absolute number of each cell population was calculated.

## 3 Results

### 3.1 Five rounds of biopanning was used to enrich the bound phages

T7 bacteriophages expressing the cDNA library of *W. bancrofti* microfilariae started forming plaques within 6-8 hours of culture. The phages that bound to the pooled IgE mAbs were collected after each biopanning round and were PCR amplified. After each round, the number of phages that bound to the IgE mAbs increased. After five (5) rounds of biopanning, we obtained 2 x 10^-3^ phages that specifically bound to the pooled IgE mAbs **(Table 1).** After *in vitro* packaging, we calculated the number of independent clones based on the plaque-forming unit (PFU). These studies showed that the prepared T7 phage display expression library of *W. bancrofti* microfilaria contained 2 x 10^8^ independent clone. We then PCR amplified the inserts from randomly selected clones and found that the library contained >90% recombinants with average insert size of >500bp.

**Table 1.**
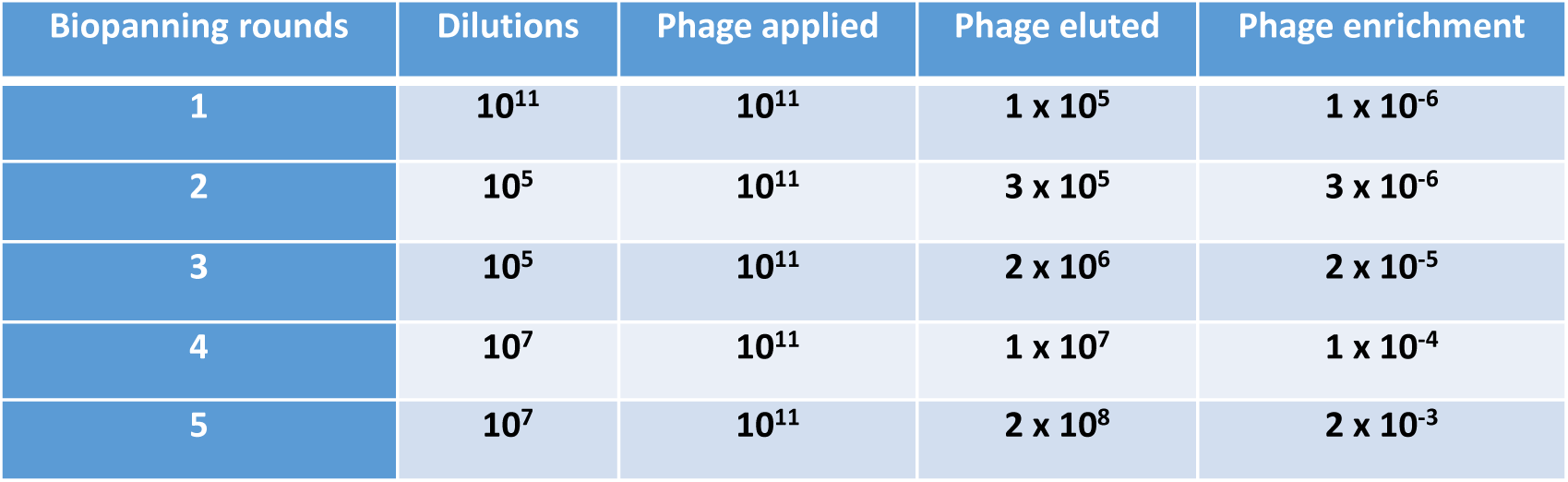
Phage enrichment of Wb Mf cDNA phage library after five rounds of biopanning.

### 3.2 IgE specific phage clones bound only to TPE sera, MF sera and pooled IgE

We then wanted to check if *W. bancrofti* infected subjects have circulating antibodies against Bm2855. In these studies, we performed the plaque assay using sera samples (i) from *W. bancrofti* infected patients showing clinical symptoms of TPE, (ii) from infected subjects with circulating microfilariae in their blood (MF), or (iii) from individuals who are naturally immune to the LF infection (EN). Pooled IgE mAbs served as the positive control. These studies confirmed that the clones bound only to TPE, MF and pooled IgE mAbs, but not to the EN sera (data not shown). We then selected 15 clones from those that bound to the pooled IgE mAbs, PCR amplified the inserts and the PCR products were separated on a 1% agarose gel. These analyses showed that all 15 clones selected had inserts with size ranging between 250bp-350bp (**Figure 1**).

**Figure 1.**
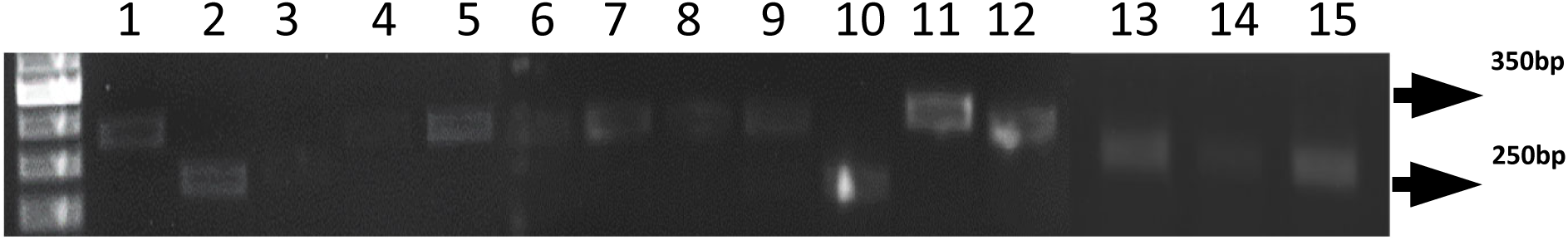
Identification of *Wb* Mf clones by PCR. 15 bacteriophage clones from those that bound to the pooled IgE mAbs were randomly picked and the DNA insert in each of the clones were PCR amplified with T7 primers. The PCR products obtained were separated on a 1% agarose gel. Our results showed that all the 15 clones had inserts and the insert sizes ranged between 250 bp-350 bp.

### 3.3 Sequence analysis identified a novel pan allergen

PCR amplified inserts were then send to the University of Illinois DNA sequencing facility for Sanger sequencing. All inserts had DNA. The nucleotide sequences obtained from the PCR inserts were then BLAST analyzed at the NCBI site (https://www.ncbi.nlm.nih.gov/gene). Out of the 15 clones, 11 clones had sequence similarity with *B. malayi* RNA recognition motif domain containing protein (*Bm*2855, NCBI Reference Sequence: XM_001896236.2, only partial RNA was deposited). A search showed that the *Bm*2855 gene has not been annotated.

The remaining four (4) clones showed sequence similarity with *W. bancrofti* translationally controlled tumor protein-like protein mRNA, (complete cds deposited in GenBank with accession # AY039808.1). Thus, our results showed that the pooled IgE mAbs bound to two proteins, *Bm*2855 and *Wb*TCTP.

We have previously characterized *Wb*TCTP and have demonstrated that it is a potent calcium-binding histamine releasing factor (HRF) [19]. Since we already characterized r*Wb*TCTP, we decided to focus on *Bm*2855 gene in this study. Furthermore, several recent reports from Kawakami’s group showed that human HRF binds with high affinity to human IgE from allergic patients [23–25]. Therefore, we decided not to analyze the *Wb*TCTP further.

We performed a phylogenetic tree analysis on the *Bm*2855 sequence to determine the evolutionary changes in the gene and to identify any similarities among different species of nematode parasites including *W. bancrofti* (**Figure 2**). Since Bm2855 binds to IgE antibodies that are isolated from subjects showing TPE clinical symptoms, we believe that Bm2855 is an allergen and possibly has a significant role in the TPE pathology. The fact that Bm2855 homologous proteins are present in the genome of several nematode parasites, we believe that this family of proteins are allergens of nematodes. Hence, we named this protein as nematode pan allergen (NPA). We then PCR amplified the full DNA sequence of NPA from *W. bancrofti* microfilaria cDNA library using sequence specific forward and reverse primers. This full-length DNA sequence of *Wb*NPA is 522 bp. The *W. bancrofti* NPA sequence is now deposited at the NCBI GenBank (Accession # ON023112.1).

**Figure 2.**
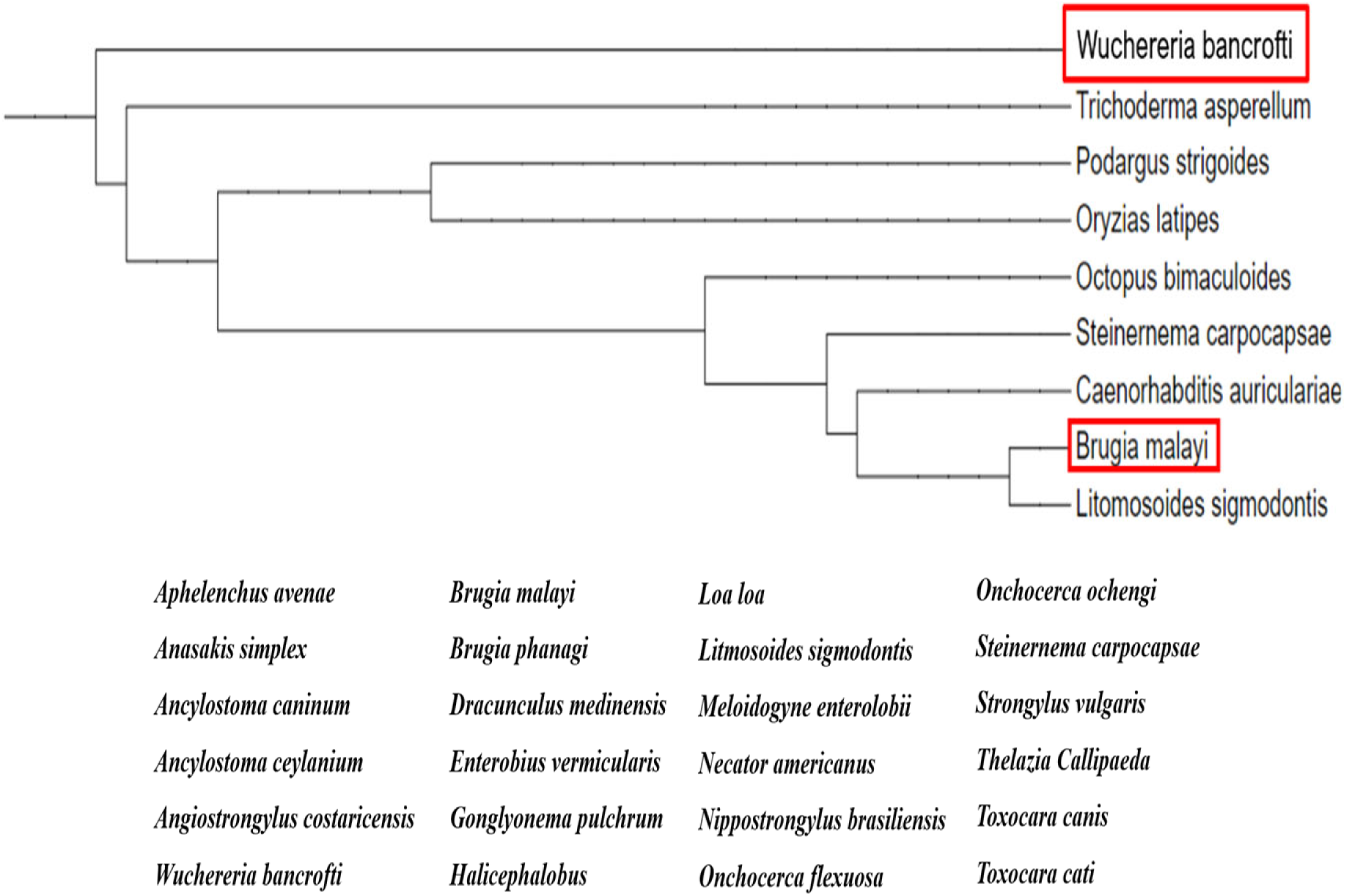
Phylogenetic tree analysis. The PCR products of the DNA insert from the bacteriophages were sequenced at the UIC Sanger sequencing facility. The nucleotide sequences obtained from all 15 clones were subjected to NCBI BLAST analysis. This identified two proteins, Bm2855 (GenBank, Gene ID 6099714), which was not annotated previously and *W. bancrofti* Translationally Controlled Tumor Protein (TCTP), which was previously described from our laboratory [30]. Based on the CLUSTAL Omega multiple sequence alignment data of Bm2855, we constructed a phylogenetic tree. Our results showed that Bm2855 of *B. malayi* and *W. bancrofti* share over 95% sequence homology. Bm2855 homologue protein were also present in several nematodes. However, there is no Bm2855 homologous protein in the humans. Given the significant homology in several nematodes and the fact that the Bm2855 protein is recognized by IgE antibodies, the sequence was named as Nematode Pan Allergen (NPA) and deposited in the NCBI GenBank with accession number ON023112.

### 3.4 The *Wb*NPA gene is expressed in different life-cycle stages of *B. malayi*

We then PCR amplified the *Wb*NPA gene from the cDNA libraries of different life cycle stages of *W bancrofti* and found that the majority of *Wb*NPA transcripts are present in the microfilarial stage, followed by the adult female worms (**Figure 3A**). Low levels of *Wb*NPA transcripts were detectable in L3 stages (**Figure 3B**).

**Figure 3.**
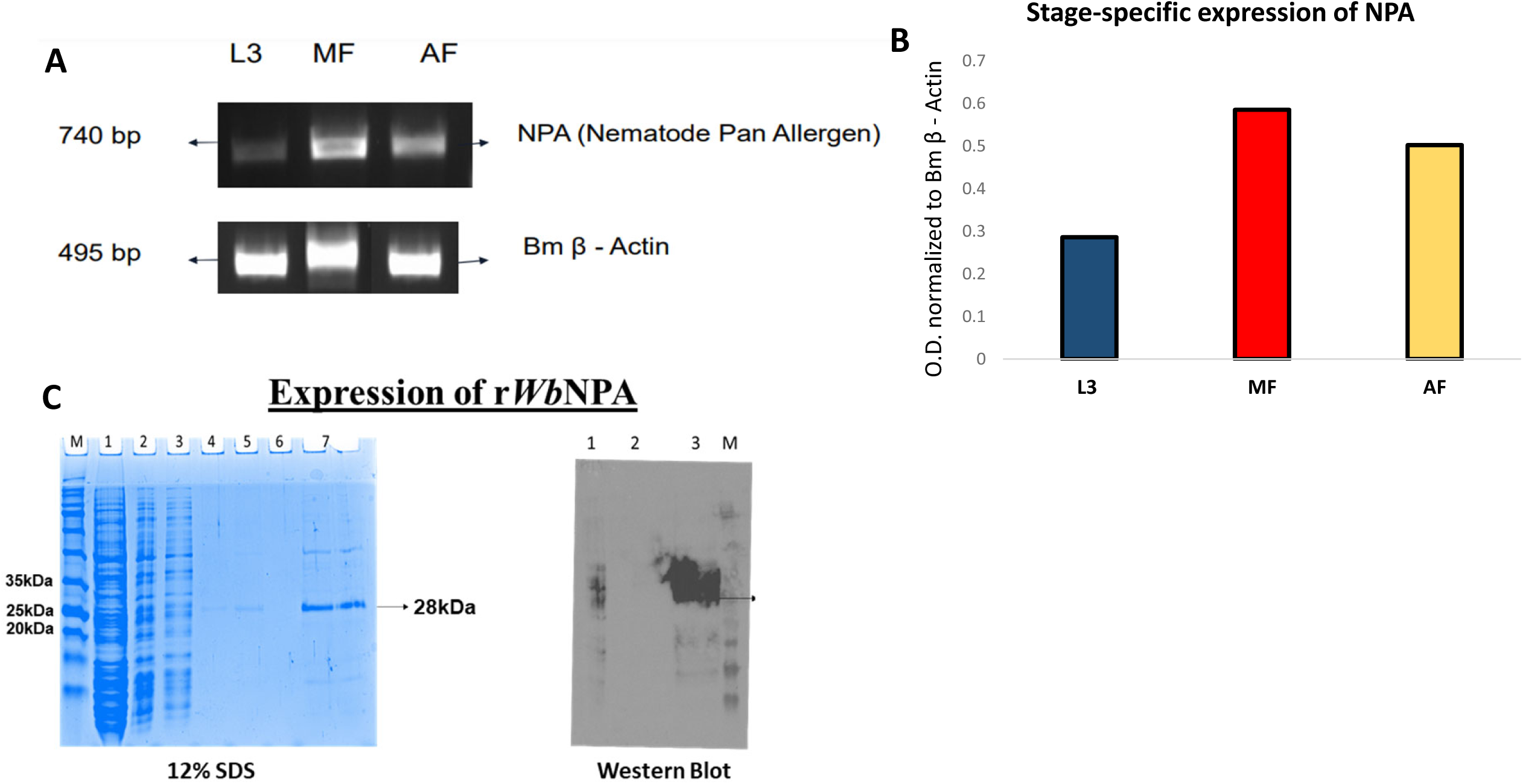
Stage Specific Expression of WbNPA in *W. bancrofti*. WbNPA gene was PCR amplified using WbNPA sequence-specific primers from the various cDNA libraries of *W. bancrofti* life cycle stages. Mf – microfilarial stage; AF – Adult female stage; L3 – Infective stage larvae. The PCR products were separated on 1% agarose gel and visualized. The intensity of the bands was normalized with Bm β-actin housekeeping gene using ImageJ software. Figure 3B: High levels of WbNPA transcript was present in Mf and AF cDNA libraries. Low levels of WbNPA was present in L3 cDNA library compared to β-actin controls. Figure 3 **C:** rWbNPA and rWbTCTP proteins were expressed and purified. WbNPA were cloned in a pREST A vector and expressed as his-tag fusion protein in BL21 DE3* *E. coli* competent cells. The crude proteins were purified, desalted, and concentrated. The purified rWbNPA protein had an estimated molecular mass of 28 kDa on a 12% SDS-PAGE gel and confirmed on a Western blot with a mouse monoclonal HRP labelled anti-His tag antibodies.

### 3.5 r*Wb*NPA was expressed and purified

*Wb*NPA cloned in a pREST A vector was expressed as his-tag fusion protein in BL21 DE3* *E.coli* competent cells, the crude protein was purified, desalted, and concentrated in endotoxin free media. The purified concentrated proteins had a molecular mass of 28 kDa (**Figure 3C**). After passing the final preparation through the endotoxin removal column, both r*Wb*NPA and r*Wb*TCTP had no detectable traces of endotoxin. Detection limit of the assay was (>0.01EU/ml).

### 3.6 r*Wb*NPA bound to a single human IgE mAb

We performed an ELISA to measure the binding of r*Wb*NPA to each of the 15 human IgE mAbs. Our results showed that the r*Wb*NPA bound to one of the monoclonal IgE antibodies (5D2) (**Figure 4**).

**Figure 4.**
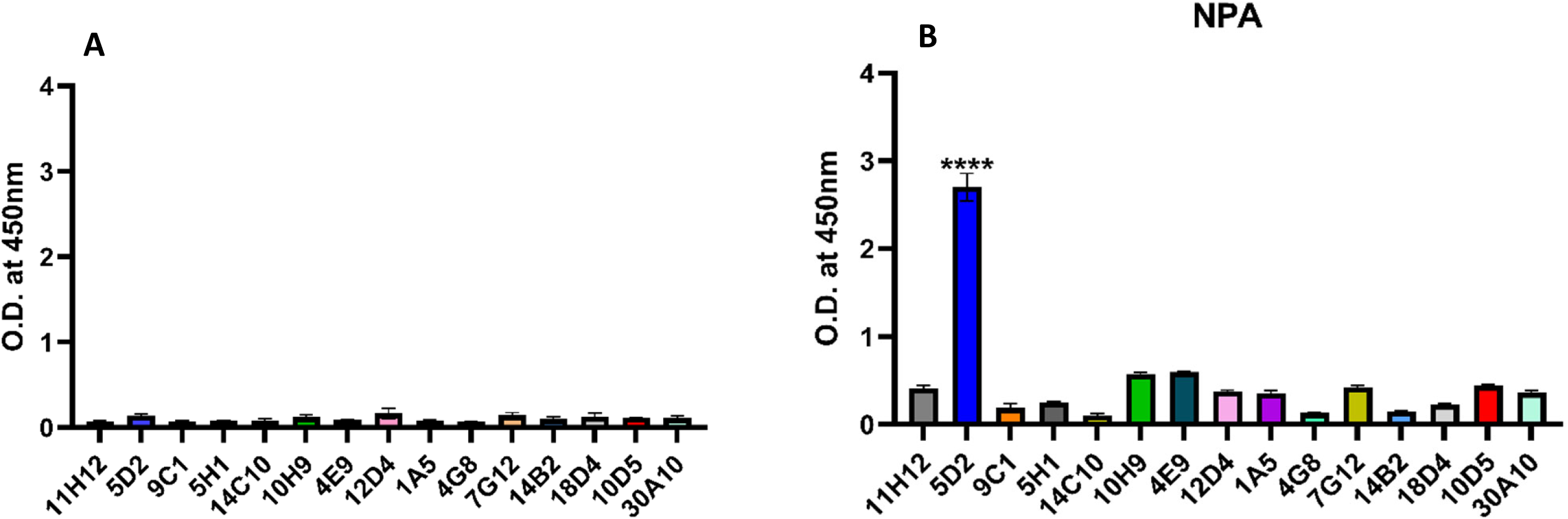
rWbNPA bound to a single IgE mAb. An indirect ELISA was performed to determine which of the 15 IgE mAbs binds to rWbNPA protein. Briefly, 96 well ELISA plates were coated with the respective proteins, after blocking, each of the monoclonal IgE antibodies were added to separate wells and HRP conjugated goat anti-human IgE antibodies were used as the secondary antibodies and color developed with 1-step TMB ELISA substrate. The optical density was read at OD450 nm. Our data showed that only one IgE mAb (5D2) specifically bound to both rWbNPA with higher affinity. None of the IgE mAbs reacted with a non-specific recombinant protein (rDmCol-4). The experiment was repeated twice with similar results. Data was analyzed using Brown-Forsythe and Welch ANOVA test. WbNPA binding to 5D2 was significant ****p<0.0001 compared to control wells.

### 3.7 *Wb*NPA induced release of histamine from mice peritoneal cells

Peritoneal cells were collected from four mice by peritoneal wash, counted the mast cells and re- suspend the cells in culture media. About one µg of r*Wb*NPA and 5D2 IgE mAb were added to the peritoneal cells and incubated for 30 min at 37^0^C. The supernatants were collected 24 hrs later, and the levels of histamine was measured. Our data showed that significant levels of histamine was released when r*Wb*NPA was added to the cells compared to the controls (**Figure 5**). In another experiment, we also added r*Wb*NPA alone to the peritoneal cells without 5D2 mAb. Our results show that the r*Wb*NPA can release histamine from peritoneal cells in the absence of IgE antibodies as well (data not shown).

**Figure 5.**
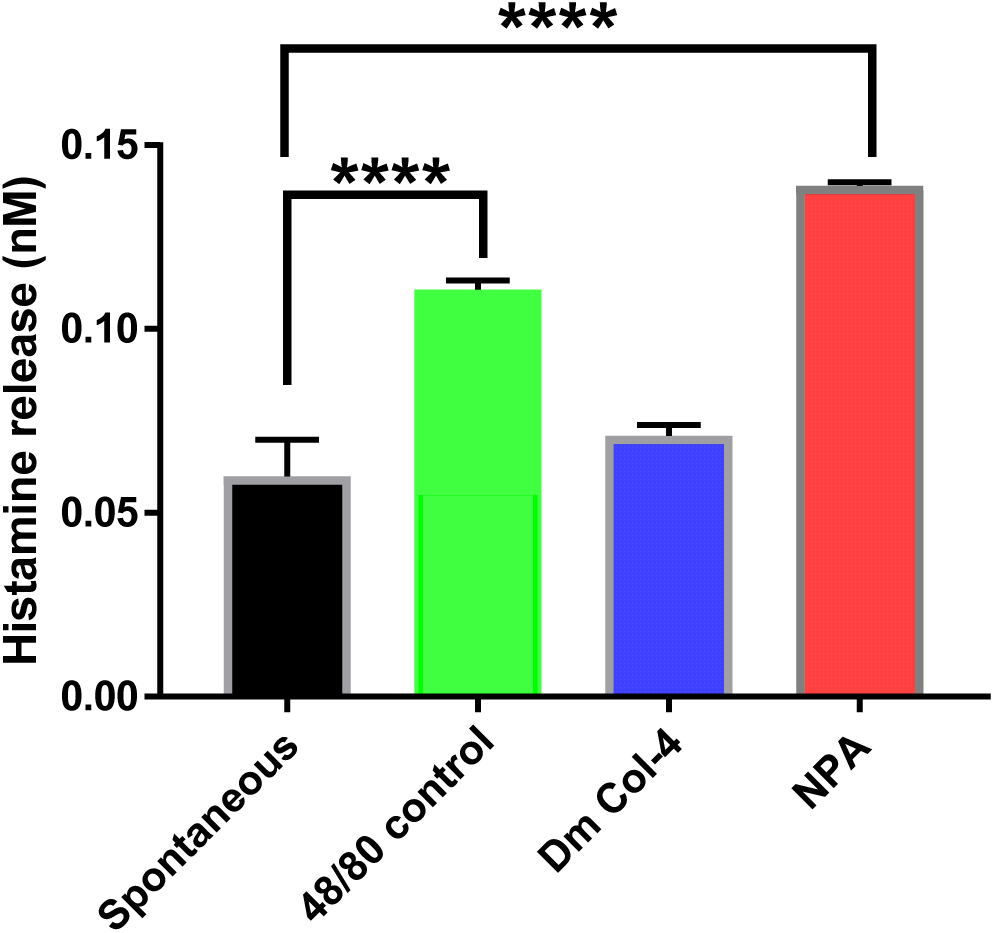
rWbNPA is a potent histamine releasing protein. Mouse peritoneal cells containing approximately 1×10^5^ mast cells were incubated with one µg/ml of rWbNPA for 30 mins at 37°C. Following incubation, levels of histamine in the culture supernatants was determined using a calorimetric histamine release assay. Substance 48/80 and rDmCol-4 was used as positive and negative controls respectively. Data obtained were analyzed using a one-way ANOVA. Our results show that rWbNPA can induce significant histamine release comparable to or higher than the substance 48/80 positive control. Significance ****p<0.0001 and n=3.

### 3.8 r*Wb*NPA can induce the release of inflammatory cytokines from mouse alveolar macrophages

Mouse alveolar macrophages collected by lavaging the alveolar spaces were incubated with 1 µg of r*Wb*NPA for 48 hrs at 37^0^C. Levels of various cytokines in the culture supernatant were measured using a BD cytokine array. Our results show that the r*Wb*NPA induced the release of the proinflammatory cytokines, IL-6 and TNF-α from alveolar macrophages (**Figure 6**).

**Figure 6.**
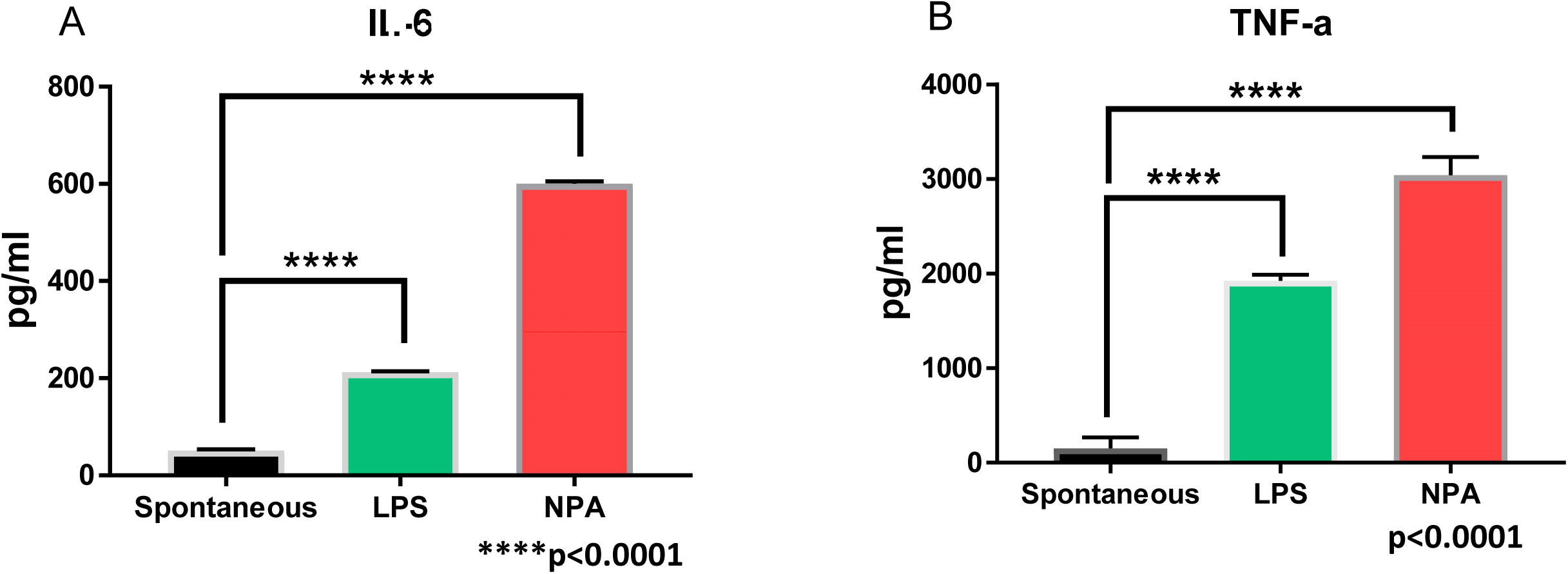
Mouse alveolar macrophages release pro-inflammatory cytokines when exposed to rWbNPA. Approximately, 2×10^4^ alveolar macrophages/well collected by bronchoalveolar lavage was seeded on to a 48 well plate. After the cells reached 70% confluence about 1µg of endotoxin free rWbNPA was added to the cells, and incubated at 37^0^C and 5% CO_2_. Following incubation, the supernatant was collected at 72hrs and analyzed for inflammatory cytokines using mouse Th1/Th2/Th17 Cytometric Bead Array in a BD flow cytometer. Our results showed an increase in the levels of TNF-α and IL-6 in cultures treated with rWbNPA compared to controls. Samples were analyzed using a one-way ANOVA. Significant ****p<0.0001 compared to controls. n=3.

### 3.9 Cellular response to *Wb*NPA

To determine if r*Wb*NPA can induce eosinophil migration, we injected 5 µg of r*Wb*NPA into the peritoneal cavity of ovalbumin-sensitized mice, and about 24 hrs later, accumulated cells were collected by peritoneal lavage and differential cell count performed. Our results show that the absolute numbers of eosinophils and neutrophils were significantly increased in the peritoneal cavity of mice injected with r*Wb*NPA **(Figure 7).** There was a slight decrease in the absolute counts of macrophages, lymphocytes, and mast cells.

**Figure 7.**
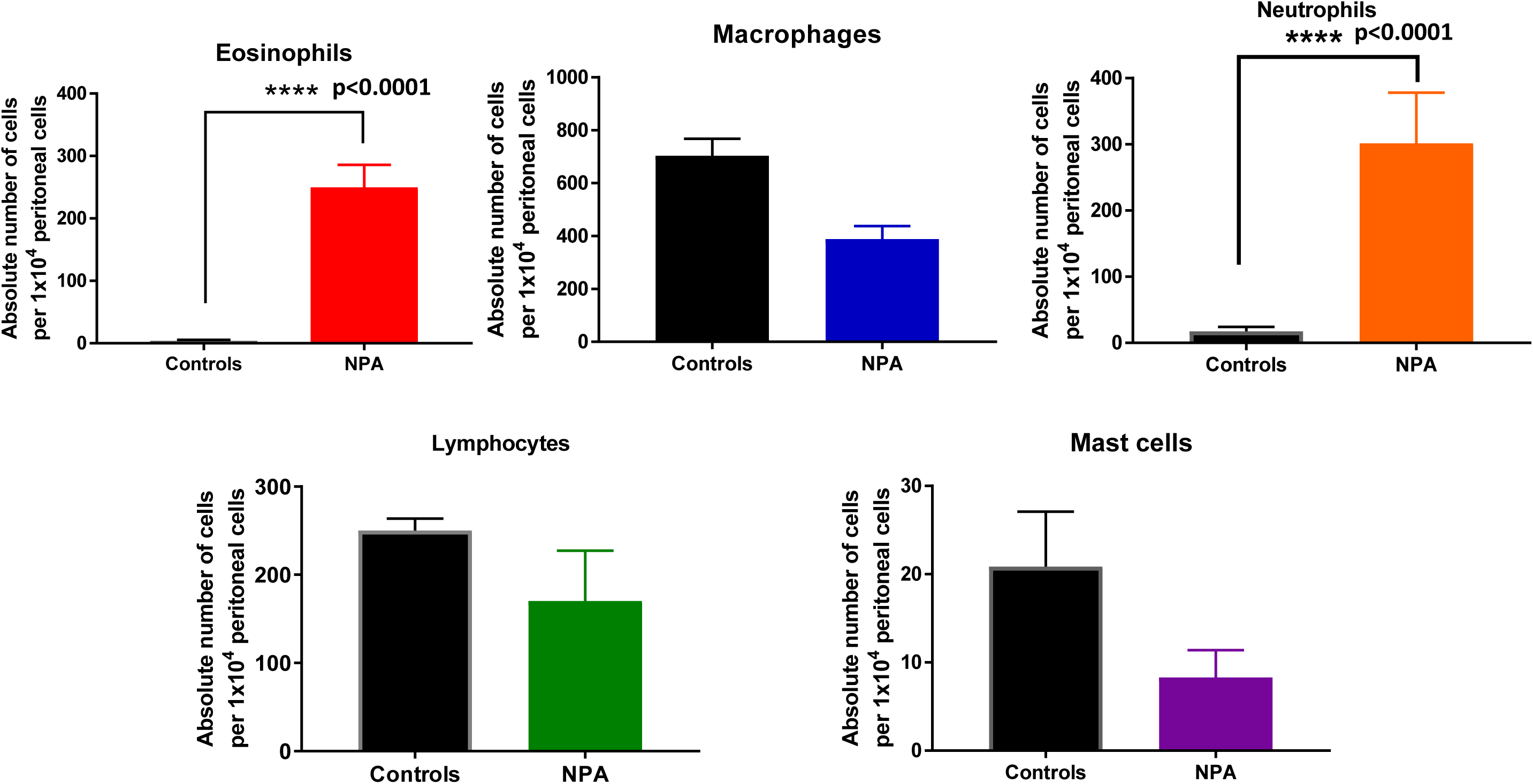
rWbNPA induced eosinophil accumulation in the peritoneal cavity. To determine if rWbNPA can induce eosinophil migration into the peritoneal cavity of mice, we injected rWbNPA into the peritoneal cavity of ovalbumin-sensitized mice. After 24 hrs, the accumulated cells were collected by peritoneal lavage and differential cell count performed. Our results show that the absolute numbers of eosinophils and neutrophils were significantly increased in the peritoneum of mice injected with rWbNPA. However, the absolute counts of macrophages, lymphocytes, and mast cells were slightly decreased. Significance ****p<0.0001, average of 10 random fields at 400x.

**Figure 8.**
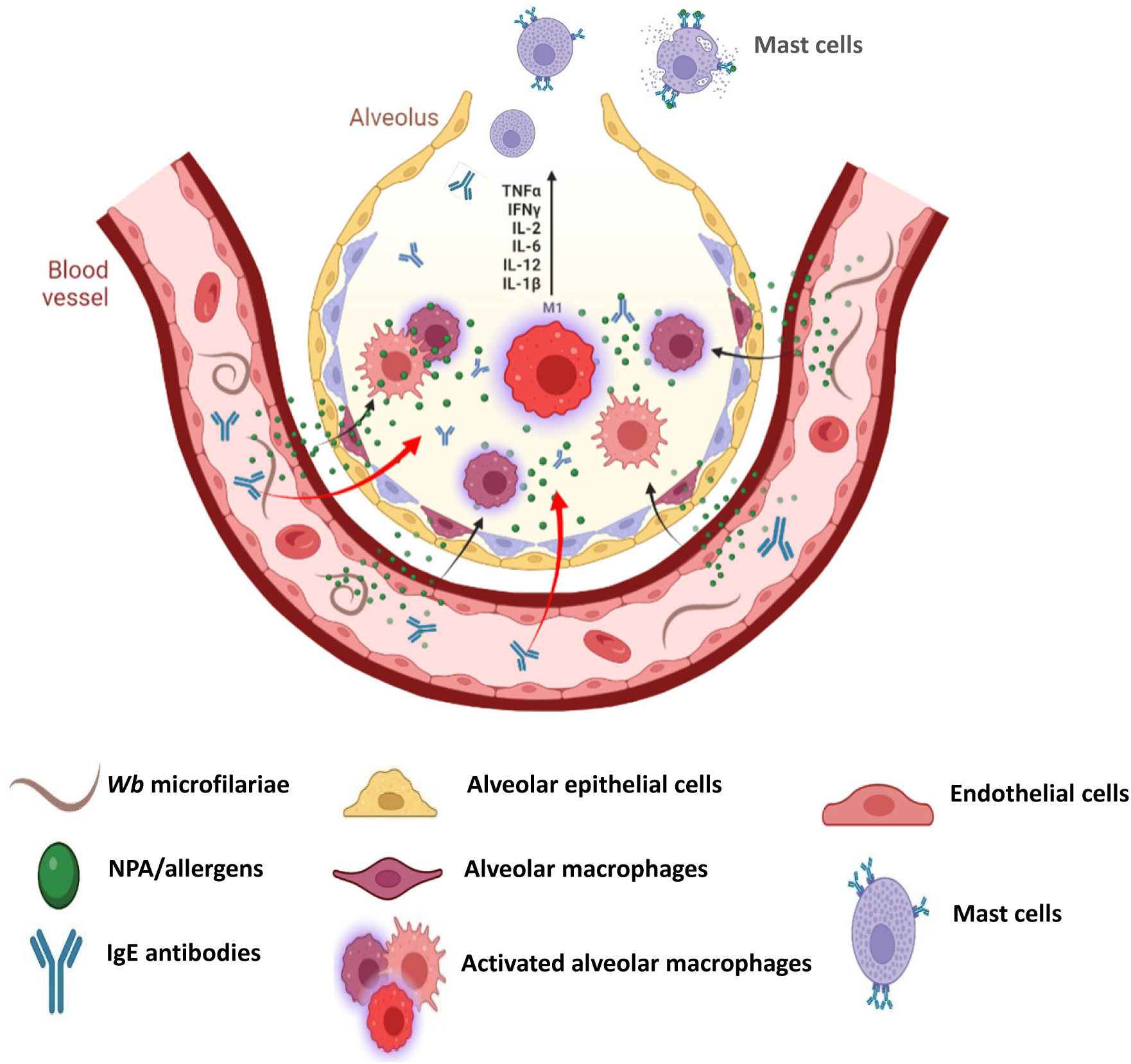
Schematic diagram of potential WbNPA role in TPE. The blood vessels surrounding the alveolus has a thin layer of endothelial cells that are close to the alveolar epithelial cells, potentially allowing easy transfer of WbNPA and IgE antibodies into the alveolus. The WbNPA within the alveolus can initiate the activation of alveolar macrophages leading to the production of proinflammatory cytokines, and initiating the inflammatory immune response. The IgE antibodies can bind to the FcεRI on the surface of mast cells. FcεRI crosslinking by WbNPA can trigger the release of histamine and other molecules by mast cells. WbNPA can also independently trigger the accumulation of eosinophils in the pulmonary tissue potentially leading to TPE clinical symptoms.

## 4 Discussion

Tropical Pulmonary Eosinophilia (TPE) is an occult manifestation of lymphatic filariasis, characterized by an allergic reaction in the respiratory tract with accumulation of eosinophils [7–9]. Allergens released from the microfilarial stages of the parasite are thought to play a significant role in the cause of TPE [26–28]. However, the allergens that trigger the TPE reaction is not known. In this study using human monoclonal IgE antibodies isolated from filarial infected subjects, we identified a 28 kDa nematode pan allergen from *W. bancrofti* (*Wb*NPA) and demonstrated a potential pathogenic role in TPE. Our studies showed that the r*Wb*NPA can release histamine from mast cells, release proinflammatory cytokines from alveolar macrophages and can promote accumulation of eosinophils in the peritoneal cavity. The human monoclonal IgE antibodies also identified a 27 kDa *W. bancrofti* translationally controlled tumor protein (*Wb*TCTP) as another allergen that may also have a potential role in TPE. In addition, our studies showed that homologous of NPA genes are expressed in several nematode parasites, making this molecule a pan allergen of nematode parasites.

Lobos and coworkers were the first to demonstrate the presence of a 23-25 kDa allergen, Bm2325 in the whole worm homogenate of *B. malayi* that reacted with TPE sera [29]. Subsequently they characterized the Bm2325 as γ-glutamyl transpeptidase (γ-GT), a key enzyme in the synthesis and degradation of glutathione [14] and might participate in pulmonary inflammation [30]. Previously our laboratory used a T7 phage-display cDNA expression library to screen the sera of lymphatic filariasis patients to identify vaccine candidates for human lymphatic filariasis [17]. In this study, we used a similar approach to screen the T7 phage displayed WbMf cDNA expression library with 15 different monoclonal IgE antibodies prepared from subjects with filarial infections, some showing TPE symptoms [16]. The human IgE monoclonal antibodies were prepared as described previously [16]. Thus, the purpose of this screening was to identify the antigens that are responsible for generating the IgE antibodies in TPE subjects. We chose Mf cDNA library because TPE clinical symptoms occur when patients have elevated levels of circulating microfilariae [1, 26]. Our transcript analysis confirmed that the *Wb*NPA gene is expressed only in the microfilariae stage and in the adult females, probably due to the presence of microfilariae in the uterus of gravid female worms [7].

Our phylogenetic studies and BLAST analysis identified the presence of NPA protein in over 25 different nematode parasites. Cross-reactive antigens have been reported between microfilaria and other parasitic nematode infections [27]. This is the first time anybody have identified a conserved pan allergen in several nematode parasites. The NPA gene have not been annotated previously from any nematode parasites, and this this is the first report on a pan allergen. IgE, eosinophils and allergy are hallmarks of nematode parasitic infections and TPE [31]. We believe that the NPA could play a major role in triggering the allergic responses in the host during parasitic larval invasion.

One of our hypothesis was that the immune response to antigens released from the circulating microfilariae results in the production of IgE antibodies (**Figure 10**). These IgE antibodies bind to mast cells lining the upper respiratory tract and cause release of histamine that promote the allergic response in the lung and respiratory tract, leading to the typical clinical symptoms of TPE [4, 9, 11, 13, 24]. In this study, we demonstrated that the r*Wb*NPA could induce the release of significant levels of histamine from mast cells in an IgE-dependent fashion. Out of the 15 IgE mAbs that we screened, only one IgE mAb (5D2) bound to r*Wb*NPA. Thus, based on these findings, we conclude that NPA produced from microfilariae can cross the endothelial barrier and enter the alveolar space (**Figure 10**). Within the alveoli, we believe that the NPA can activate alveolar macrophages to release proinflammatory cytokines. A role for lung macrophages in TPE have been reported by other investigators as well [11, 32]. A report by Maya and coworkers [33] demonstrated that the saline extract of microfilariae in low concentrations could activate rat lung cells *in vitro*. Thus, the microfilarial antigens can directly activate alveolar macrophages.

Another clinical finding in TPE is the accumulation of eosinophils in the lung [4, 10, 11, 32]. Although we did not evaluate the eosinophil response to WbNPA in the lung tissue, our studies demonstrate that injection of WbNPA into the peritoneal cavity of mice resulted in the recruitment of a higher number of eosinophils suggesting that WbNPA released from the microfilariae may play a significant role in recruiting eosinophils into the lungs in TPE.

In conclusion, in this study we identified a novel molecule from *W. bancrofti* microfilariae that bound to a monoclonal IgE antibodies prepared from filaria infected human subjects. Subsequent analysis showed that this molecule is an allergen and homologous gene is present in a number of parasitic nematodes. Hence, we named this molecule as Nematode Pan Allergen (NPA).

## 6 Author Contributions

SK, GM, RK, and AB planned the experiments. SK and GM performed the phage display screening, AH and SS prepared the IgE mAbs, SK and AB performed all the ELISA and histamine assays, SK and RK performed the cellular studies. SK and RK drafted the manuscript.

## 8 Acknowledgments

We would like to thank Dr. Steven A. Williams, Smith College, Northampton, MA, for supplying us the cDNA libraries of various stages of the *W. bancrofti* parasite.

